# Multi-parametric imaging of spatio-temporal cAMP signaling, transmembrane potential, and intracellular calcium in the Langendorff-perfused heart

**DOI:** 10.1101/2025.04.30.650917

**Authors:** I-Ju E. Lee, Jessica L. Caldwell, Lilian R. Mott, Lianguo Wang, Crystal M. Ripplinger

**Author notes:** **Corresponding author:** Crystal M. Ripplinger, PhD Department of Pharmacology, University of California Davis, One Shields Avenue, Davis, CA 95616.

## Abstract

The spatiotemporal dynamics of intracellular second messengers and signaling molecules, including cyclic adenosine monophosphate (cAMP), have been studied extensively in isolated cardiomyocytes using Förster resonance energy transfer (FRET)-based reporters. However, little is known about how molecular signaling events affect tissue-level function. Optical mapping of transmembrane potential (V_m_) and intracellular calcium (Ca^2+^_i_) is frequently performed in isolated intact hearts to reveal tissue-level electrophysiological function but cannot reveal molecular underpinnings. Here, we developed a novel multi-parametric optical imaging system that enables multi-parametric recording of four wavelengths to visualize cAMP and real-time electrophysiological responses (V_m_ and Ca^2+^_i_) by utilizing concurrent <u>FRE</u>T imaging and <u>d</u>ual <u>o</u>ptical <u>m</u>apping (FRE-DOM) in the intact heart from a cardiac-specific FRET-based cAMP reporter mouse. We showed that cAMP is strongly and heterogeneously activated throughout the heart (atria and ventricles) in response to β-AR stimulation, the time course of which matches heart rate, action potential, and Ca^2+^ responses. This novel imaging system will provide insight into the relationship between cAMP signaling, electrophysiology, and arrhythmogenesis at high spatio-temporal resolution.

## Introduction

The development of genetically encoded Förster resonance energy transfer (FRET)-based biosensors have been used to understand the spatiotemporal dynamics of second messengers and signaling molecules in live cells. FRET is a non-invasive technique that does not require the destruction of cells, allowing for continuous monitoring and preserving the physiological context. A number of biosensors have been created to measure cyclic nucleotide generation or sub-cellular localization (e.g., cyclic adenosine monophosphate [cAMP], cyclic guanosine monophosphate [cGMP])^1-4^, kinase activity (e.g., protein kinase A [PKA], Ca^2+^ calmodulin kinase II [CaMKII])^5-7^, and associations between the sarcoplasmic reticulum ATPase (SERCA) pump and phospholamban^8^ in live cardiomyocytes. While FRET sensors offer valuable insights into sub-cellular compartmentalization, target activity, and nanodomain signaling, isolated cardiomyocytes have limitations when removed from the intact heart. For example, it is unclear if certain electrophysiological events, including early or delayed afterdepolarizations (EADs, DADs) in an individual myocyte will overcome the source-sink mismatch to trigger premature ventricular complexes (PVCs) in tissue^9^. Moreover, isolated cardiomyocytes are often pooled from throughout the heart (e.g., left and right ventricular myocytes are combined), and the expression of ion channels and other signaling proteins can change significantly if cells are cultured^10^. These approaches also fail to account for intrinsic regional heterogeneity within the myocardium, as gradients in ion channel expression and sympathetic innervation, such as those between the ventricular base and apex, are lost when cells are dissociated and studied in isolation^11^.

Cardiac optical mapping is a fluorescence-based technique which offers unprecedented spatial resolution and real-time visualization of electrical activity across the surface of intact Langendorff-perfused hearts^12^. This method allows for direct, contactless recording of transmembrane potential (V_m_) and optical action potentials, facilitating the measurement of electrical activity and the identification of tissue heterogeneities that can lead to arrhythmias^13^. Additionally, optical mapping can directly image intracellular calcium (Ca^2+^_i_) transients, capturing the rise and fall of Ca^2+^_i_ that triggers the contraction and relaxation of cardiac muscle cells^14^. Despite these advantages, cardiac optical mapping gives little insight into the underlying molecular mechanisms. To gain cellular and molecular insight, recent advances in tissue clearing have allowed for labeling of proteins of interest in intact hearts^15,16^, but relating real-time molecular signaling events to whole-heart function has not been performed.

To address this gap, we initially chose to image cAMP at the whole-heart level because it is a crucial second messenger in the adrenergic signaling cascade, and there is limited understanding of macroscale heterogeneity of cAMP signaling throughout the heart and its contribution to arrhythmogenesis. Here, we generated a cardiac-specific *CAMPER* reporter mouse^17^ expressing a FRET-based Epac-mediated cAMP biosensor (termed *CAMPER* mouse) and developed a novel integrated whole-heart optical imaging system capable of simultaneous <u>FRE</u>T imaging of cAMP activity along with <u>d</u>ual <u>o</u>ptical <u>m</u>apping of V_m_ and Ca^2+^_i_ (FRE-DOM). This approach uses cAMP as a readout of sympathetic activity and V_m_ and Ca^2+^_i_ as a measure of downstream electrophysiological function. To our knowledge, this is the first multi-parametric imaging system capable of assessing functional electrophysiological outputs in response to cellular signaling events at whole-heart level following β-adrenergic receptor ((β-AR) stimulation. We also show real-time cAMP and V_m_ activation in the atria and locate the earliest activation point, allowing us to determine the cAMP response in sinoatrial nodal (SAN) tissues. To test the heart’s response to physiological nerve activity, we demonstrate the spatio-temporal kinetics of cAMP generation in response to chemically-induced sympathetic nerve stimulation, which is known to have important functional differences compared to *in vitro* agonist application^18^. Overall, our multiparametric imaging system provides a powerful tool to study the spatial heterogeneity of cAMP activity and its electrophysiological responses, offering new insights into the mechanisms that may underlie arrhythmia susceptibility in the diseased heart.

## Materials and Methods

### Cardiac-specific *CAMPER* Mice

Generation of cardiac-specific *CAMPER* mice was described previously^19^. In brief, the *CAMPER* floxed mouse (Jackson Laboratories #032205), that reports cAMP binding by changes in FRET between donor (CFP) and acceptor (YFP) at the cAMP binding domain of Epac1^20^, was crossed with the cardiac-specific alpha myosin-heavy chain (Myh6) Cre mouse (Jackson Laboratories #011038) for cardiac-specific expression. Mice were genotyped before use, and only homozygous male and female *CAMPER* mice 17 ± 5 weeks old were used for experiments.

### Quadruple-parametric imaging system set up

A multi-parametric whole-heart imaging system was built by modifying an existing optical mapping setup (THT Mesoscope, SciMedia, Fig 1A). Two light emitting diode (LED) excitation sources at 457 nm (Blue LED: T-LED+, Sutter Instrument) and 530 nm (Green LED: LEX-2, SciMedia) with collimators were used for epi-illumination of the heart to excite the FRET sensor and exogenous dyes, respectively. The emitted light signal is collected by a Planapo 1X lens (objective lens, WD = 61.5 mm, SciMedia) with infinity correction. The lens focuses the collected light at infinity and this correction allows multiple layers of filter sets and cameras to be introduced in the light path. The different wavelengths of the emitted light were separated via dichroic mirrors and filters, as depicted in Fig 1B, and then passed through the secondary Planapo 1X lens (condensing lens, WD = 61.5 mm, SciMedia) prior to being captured by the cameras. The use of two lenses of the same focal length in tandem lens configuration results in a magnification of 1X (pixel size = 0.1 mm)^21-23^. The mapping system is equipped with one high-resolution sCMOS camera (Prime BSI, Photometrics) for FRET image acquisition and two high-speed CMOS cameras (MiCAM ULTIMA-L, SciMedia) for mapping V_m_ and Ca^2+^.

**Figure 1.**
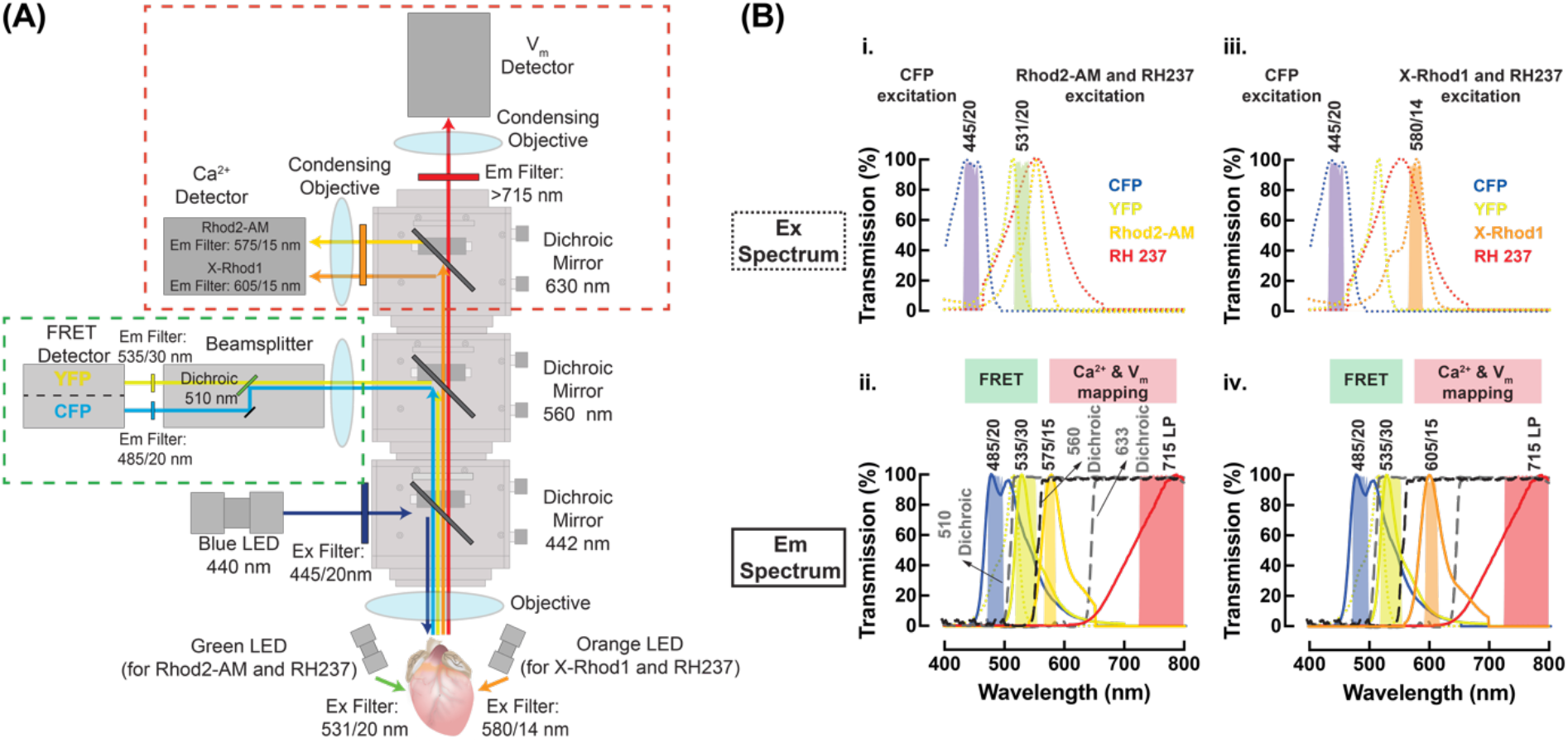
Quadruple-parametric FRET and dual optical mapping (FRE-DOM) system. **(A)** Schematic of the quadruple-parametric imaging system illustrating the optics used and the light path for cAMP FRET, V_m_ and Ca^2+^ signals. **(B**_**i**._**), (B**_**iii**._) Excitation and **(B**_**ii**._**), (B**_**iv**._) emission spectra of the three parameters – cAMP FRET (CFP and YFP), Rhod2-AM or X-Rhod1 (Ca^2+^), and RH237 (V_m_) fluorescence, illustrating the spectral separation of the three signals as implemented in this system. Dotted lines: excitation wavelength, solid lines: emission wavelengths, long-dotted vertical lines: dichroic mirrors, shaded boxes: filters.

For FRET imaging, CFP is excited at 445/20nm with the blue LED source (Fig. 1B_i._) and CFP/YFP emission is reflected by the first long-pass dichroic at 560 nm (Fig. 1B_ii._), collected by an image splitting device (OptoSplit II, Cairn Research, Ltd) and split onto a single sCMOS detector at 485/20nm (CFP) and 535/30nm (YFP) (Fig. 1B_ii._). For dual optical mapping, hearts were loaded with RH237 (V_m_) and Rhod2-AM (Ca^2+^), and excited at 531/20nm with the green LED source (Fig. 1B_i._). RH237 and Rhod2-AM emission are transmitted by the first dichroic mirror, split by a second dichroic mirror at 633nm (Fig. 1B_ii._), and collected by two high-speed CMOS cameras at 715nm/LP (RH237) and 575/15nm (Rhod2-AM) (Fig. 1B_ii._). The system also supports imaging of RH237 and the red-shifted Ca^2+^ indicator, X-Rhod1, with 580nm excitation (Fig. 1B_iii._) and emission at 605/15nm (Fig. 1B_iv._). This dye combination will allow for greater spectral separation and simultaneous (rather than interleaved) excitation of the FRET fluorophore and exogenous dyes.

At the start of each experiment, a focusing target was positioned in front of the objective lens, and its position was adjusted until all three cameras were in clear focus. VisaView software (Visitron Systems GmbH) is utilized to control the sCMOS camera configuration, OptoSplit image alignment, and the blue LED on/off triggering. For the manual alignment of OptoSplit, VisaView facilitates the process by overlaying live images with contrasting colors, which effectively highlight misalignment. This real-time feedback enables rapid alignment of images through adjustment of the control knobs on the OptoSplit. BV Workbench (Brainvision, Inc.) serves as the software interface for the two high-speed CMOS cameras and the green LED on/off triggering. The CMOS cameras were spatially aligned by utilizing the Camera Calibration function within the software. This function superimposes images from different cameras and uses an edge detection algorithm, enabling manual adjustment of the angle of the dichroic mirrors until all fields of view are spatially aligned.

### Whole-heart Langendorff perfusion

*CAMPER* mice received an intraperitoneal injection of heparin (100 IU) and were anesthetized with pentobarbital sodium (>150 mg/kg). For Langendorff perfusion, hearts were prepared according to previously established methods^24-26^. In short, hearts were excised through a mid-sternal incision, placed in cold cardioplegia solution (with composition in mmol/L: NaCl 110, CaCl_2_ 1.2, KCl 16, MgCl_2_ 16, and NaHCO_3_ 10), and then cannulated at the ascending aorta. Subsequently, the hearts were retrograde perfused through the aorta with oxygenated modified Tyrode’s solution (in mM: NaCl 128.2, CaCl_2_ 1.3, KCl 4.7, MgCl_2_ 1.05, NaH_2_PO_4_ 1.19, NaHCO_3_ 20, and glucose 11.1; pH 7.4) at 37°C. The perfusion chamber was heated, and the perfusion pressure was maintained at 80 mmHg. To minimize motion artifacts, blebbistatin (10 μM, Tocris Bioscience) was added to the perfusate^27^. Ag/AgCl needle electrodes were placed in the bath to record an electrocardiogram (ECG) similar to a lead I configuration.

### FRET imaging and dual optical mapping

Hearts were equilibrated after cannulation and perfusion for 10 min followed by staining with calcium (Rhod2-AM or X-Rhod1)- and voltage (RH237)-sensitive dyes. A 1 ml solution containing Rhod2-AM (50 μl of 1 mg/ml stock solution mixed with 50 μl Pluronic F-127 and 900 μl Tyrode’s solution) or X-Rhod1 (20 μl of 1 mg/ml stock solution mixed with 20 μl Pluronic F-127 and 960 μl Tyrode’s solution) was prepared and slowly injected into a port just proximal to the cannula with a 5 min dye washout period. Similarly, a 1 ml solution containing RH237 (5-10 μl of 5 mg/ml stock solution mixed with 970 μl Tyrode’s solution) was prepared and slowly injected into the dye port with a 5 min dye washout.

The *CAMPER* heart was then illuminated by two LED excitation light sources at wavelengths of 445/20 nm (blue) and 531/20 nm (green). While the former excites CFP, leading to YFP emission in the tissue, the latter excites both RH237 and Rhod2-AM dyes. With this optical setup, there is spectral overlap between YFP emission and the green LED excitation. Therefore, interleaved imaging was performed to avoid signal crosstalk. CFP and YFP time-lapse imaging was acquired with a 100 ms exposure time every 10 s. Immediately following each CFP/YFP frame, the sCMOS camera sent out a TTL signal with a delay time to the CMOS cameras triggering V_m_ and Ca^2+^ recordings at a sampling rate of 1 kHz for 2 s. If simultaneous FRET and V_m_/Ca^2+^ imaging is required, RH237 and X-Rhod1 can be used with LED excitation at 580/14 nm (orange). Since there is no spectral overlap between YFP emission and the orange LED excitation, FRET and V_m_/Ca^2+^ imaging can be acquired concurrently. CFP/YFP images were captured at a resolution of 1024 x 2048 pixels with a field of view of 7.24 mm x 14.5 mm, while V_m_ and Ca^2+^ images were recorded at a resolution of 100 x 100 pixels with a field of view of 10 mm x 10 mm.

Hearts were subjected to an acute bolus of norepinephrine (NE, 0.5 or 1.5 μM; Sigma-Aldrich) or tyramine (20 μM; Sigma-Aldrich) to test responsiveness to β-AR stimulation. Experiments were performed at 37°C under normal sinus rhythm, which allowed for assessments of heart rate changes and physiological activation and repolarization patterns.

### FRET and optical mapping data analysis

For whole heart FRET data analysis, a custom made MatLab (MathWorks) script was used to visualize cAMP responses. FRET was calculated as R/R_0_, where R = donor/acceptor (CFP/YFP) and R_0_ = baseline CFP/YFP, on a pixel-by-pixel basis and visualized as a ΔFRET map. ImageJ software was utilized to select regions of interest (ROIs) corresponding to the right ventricular (RV) base and left ventricular (LV) apex, with R/R_0_ calculated for each area. For each of the ROIs, average FRET ratio curves and FRET ratio responses at different time points were plotted.

For V_m_ and Ca^2+^ data analysis, Electromap software was used as described previously^28^. A binary mask of the epicardial surface was manually selected for whole-heart analysis. For post-processing of fluorescent signals, a spatial Gaussian filter (3 x 3 pixels) and a baseline drift correction (Top-Hat average) were used. Action potential (AP) duration (APD) at 80% (APD_80_) and Ca^2+^ transients duration (CaTD) at 50% (CaTD_50_) were computed as the time intervals from the activation time (time of maximum first derivative of the upstroke) to 80% of repolarization and 50% of calcium transient decay for 10 consecutive beats and averaged, respectively^29^. Following APD and CaTD calculations, whole heart APD and CaTD image matrix maps were exported from Electromap. Fiducial and anatomical markers were used to align images, and ImageJ software was used to select the same ROIs as the FRET data, representing RV base and LV apex for APD_80_ and CaTD_50_. FRET ratio data were colocalized with APD and CaTD values for each ROI to determine the relationships between cAMP kinetics and functional outputs.

### Statistical analysis

Data are presented as mean ± SEM from N animals. To test for differences in the shape of two signals, Kolmogorov-Smirnov test was used (GraphPad Prism). Data was considered significant when p<0.05.

## Results

### Validation of spectral separation in multi-parametric optical imaging

We first asked whether the exogenous voltage (RH237)- and calcium (Rhod2-AM or X-Rhod1)-sensitive dyes used for optical mapping would interfere with the cAMP FRET signal. To test for potential crosstalk between these dyes and the FRET fluorophores (CFP, YFP), experiments were performed on Langendorff-perfused hearts from cardiac-specific *CAMPER* mice. These hearts were initially loaded with a single fluorescent dye (Rhod2-AM) and then subjected to interleaved FRET and dual optical mapping during an acute bolus of norepinephrine (NE) infusion (0.5 μM). Following this, the hearts were subsequently loaded with RH237 and the same image acquisition protocol was repeated. Kolmogorov-Smirnov test was performed to compare the similarity of FRET signal trace between different dye loadings. When hearts were loaded with Rhod2-AM alone, slight bleed-through of the Rhod2-AM signal was observed in the V_m_ channel (Fig. 2A_i._), but no change was observed in FRET (R/R_0_) signal compared to hearts with no dye loading (Fig. 2B). When hearts were loaded with both Rhod2-AM and RH237, the raw intensity of RH237 was high enough to strongly overwhelm the bleed-through of the Rhod2-AM signal (Fig. 2A_ii._), and there was no significant difference in FRET signal compared to hearts with no dye loading or with Rhod2-AM only (Fig. 2B). These results confirm spectral separation between V_m_, Ca^2+^ and FRET signals, and that loading of the exogenous dyes does not interfere with the cAMP FRET signal.

**Figure 2.**
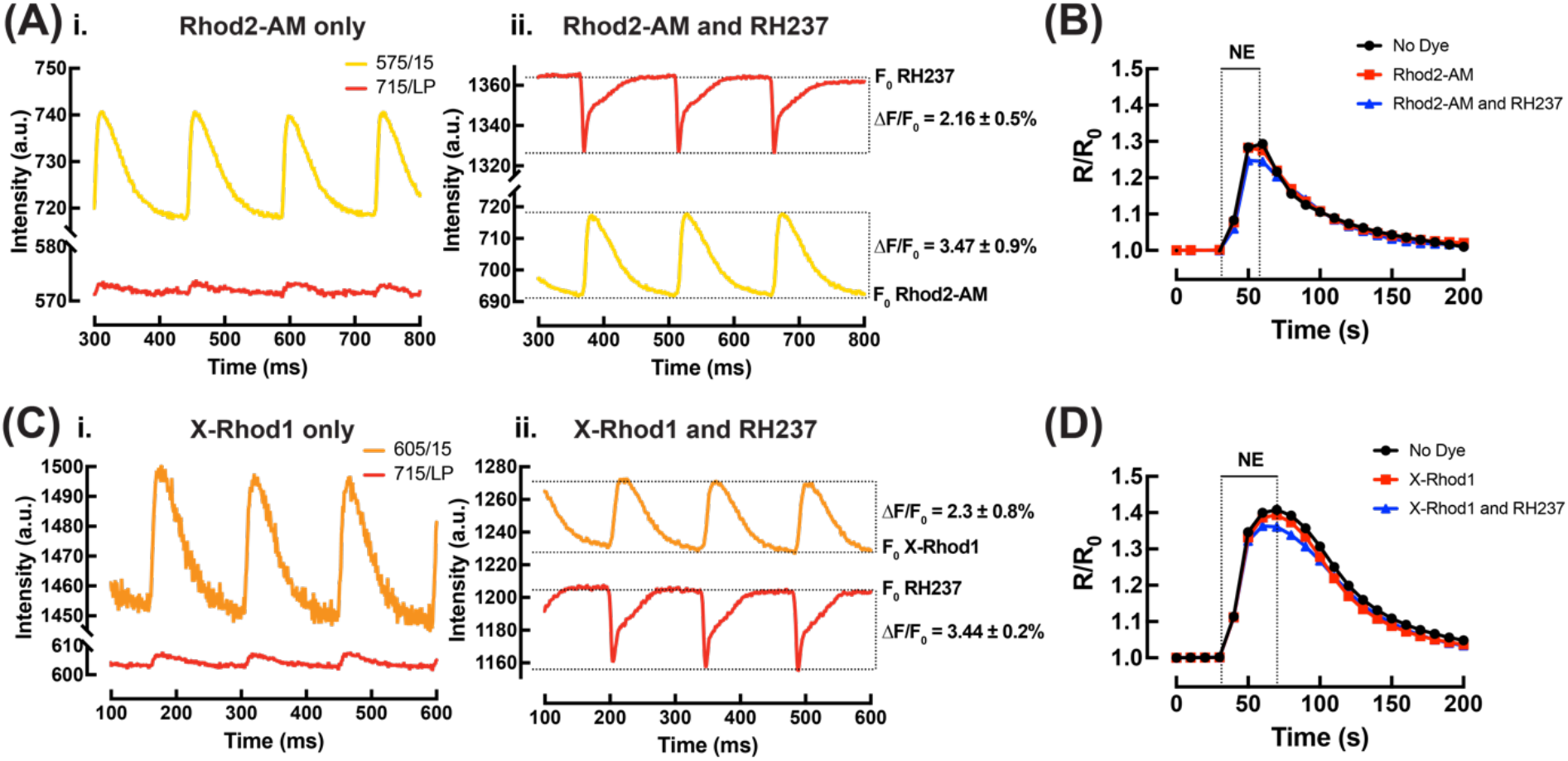
Validation of spectral separation. **(A**_**i**._) Imaging of hearts loaded with Rhod2-AM or **(C**_**i**_) X-Rhod1 only and **(A**_**ii**._**) (C**_**ii**_) following with RH237. **(B)** Representative FRET acquisition of example hearts loaded with no dyes, Rhod2-AM or **(D)** X-Rhod1 only and with RH237 during an acute bolus of 0.5 μM NE infusion. Kolmogorov-Smirnov test, No dye vs. Rhod2-AM, p = 0.3876; No dye vs. Both, p = 0.39; Rhod2-AM vs. Both, p = 0.9525. No dye vs. X-Rhod1, p > 0.99; No dye vs. Both, p = 0.9983; X-Rhod1 vs. Both, p = 0.9983, N = 1-3 hearts.

If simultaneous, rather than interleaved V_m_/Ca^2+^ and FRET imaging is desired, the red-shifted intracellular Ca^2+^ indicator (X-Rhod1) in combination with RH237 can be used. The same dye loading and imaging acquisition protocol (as above) was repeated on Langendorff-perfused *CAMPER* mouse hearts. Similar to Rhod2-AM, the X-Rhod1 signal exhibited slight bleed-through in the V_m_ channel, but this was negligible after RH237 loading (Fig. 2C_i._ and 2C_ii._). No change was observed in FRET signal with any dye loading (Fig. 2D).

### Spatio-temporal whole-heart cAMP and V_m_-Ca^2+^_i_ response following β-AR stimulation

Next, we tested the capability of our quadruple-parametric optical mapping system to simultaneously capture, cAMP (FRET (CFP/YFP)), V_m_ (RH237) and Ca^2+^_i_ (Rhod2-AM) activity during acute β-AR stimulation with a bolus of NE infusion (0.5 μM) in intact *CAMPER* hearts (N = 3). FRET/APD_80_/CaTD_50_ maps during baseline (0 sec), NE bolus (30 to 60 sec) and washout (90 to 120 sec) are shown in Fig. 3A-C. The application of an acute bolus of NE led to an increase in CFP fluorescence and a decrease in YFP fluorescence (Fig. 3D). This resulted in a significant increase in both heart rate and the FRET ratio (Fig. 3E). Similar to wild-type mouse hearts^18^, the *CAMPER* hearts displayed a biphasic APD response (Fig. 3F), where APD prolongation first occurred to maximum at ∼40 s of stimulation before shortening to near-baseline values by ∼60 s. Despite the biphasic APD response, CaTD showed a monotonic decrease in the *CAMPER* mouse hearts (Fig. 3F).

**Figure 3.**
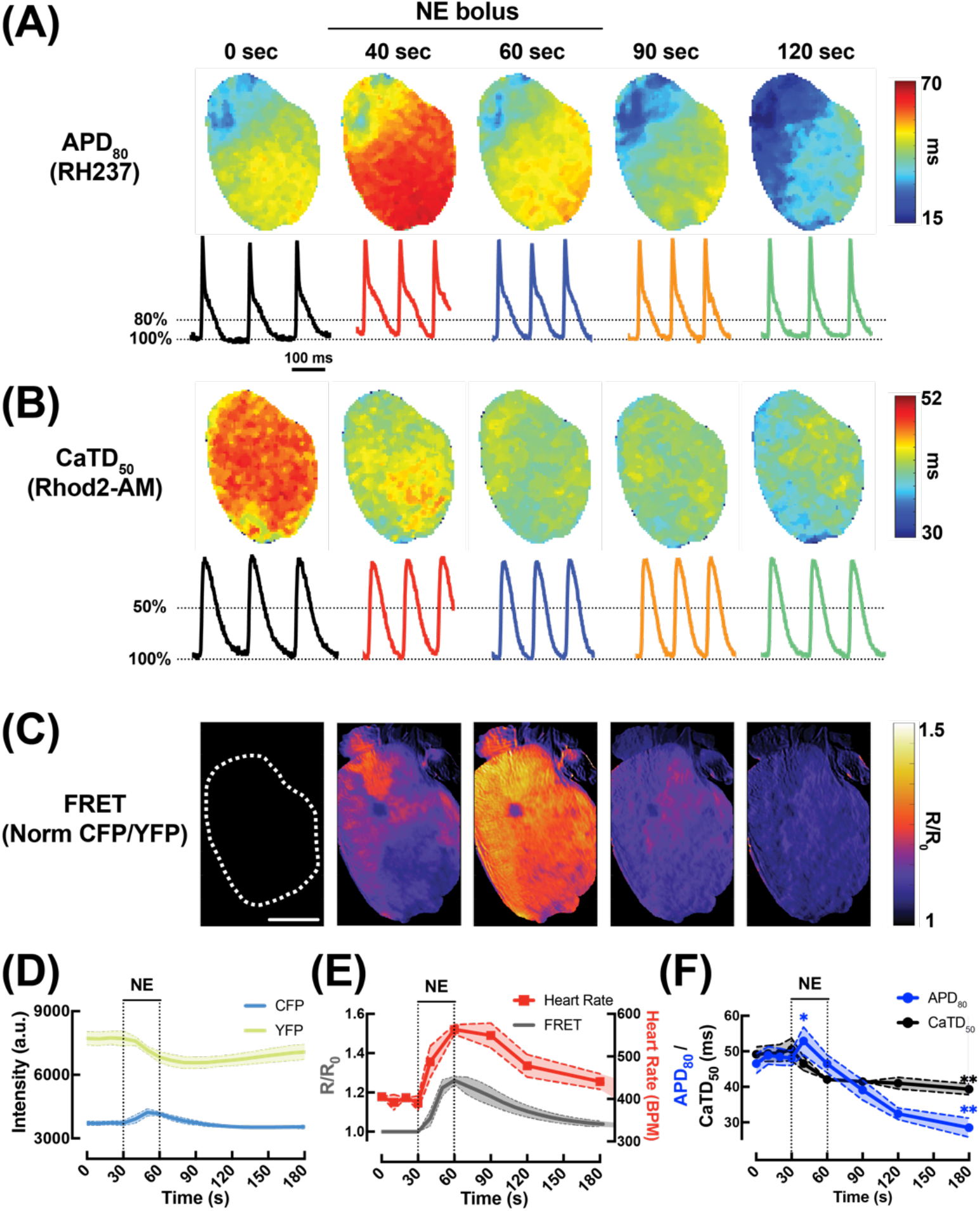
Demonstration of whole-heart cAMP and V_m_-Ca^2+^_i_ response following β-adrenergic stimulation. **(A)** Example maps of action potential duration (APD_80_), **(B)** calcium transient duration (CaTD_50_) and **(C)** pseudo-colored CFP/YFP ratio images during acute 0.5 μM NE infusion (scale bar = 3 mm). **(D)** Temporal CFP and YFP fluorescence responses in *CAMPER* mouse hearts after application of a 0.5 μM bolus NE. **(E)** Temporal CFP/YFP (normalized to baseline (R_0_)) ratio changes with heart rate and **(F)** APD_80_, CaTD_50_ changes in response to 0.5 μM NE (Paired t-test, *p < 0.05 vs. time = 0 sec; **p < 0.01 vs. time = 0 sec, N = 3 hearts).

To explore how spatio-temporal changes in cAMP signaling relate to electrophysiological responses, we analyzed two cardiac regions of interest (ROIs) from the right ventricular (RV) base and apex. An example is shown in Fig. 4. While cAMP levels followed a similar time course in both regions following β-AR stimulation, cAMP levels were modestly elevated in the RV base compared to the apex during NE bolus infusion, with the most pronounced difference observed at 40 seconds (Fig. 4B). This regional variation in cAMP activity corresponded to functional differences in CaTD_50_ and APD_80_ between the RV base and apex. Specifically, we observed a trend toward greater shortening of CaTD_50_ and greater prolongation of APD_80_ in the RV base of the heart compared with the apex at the same time point (Fig. 4C-D and 4E-F).

**Figure 4.**
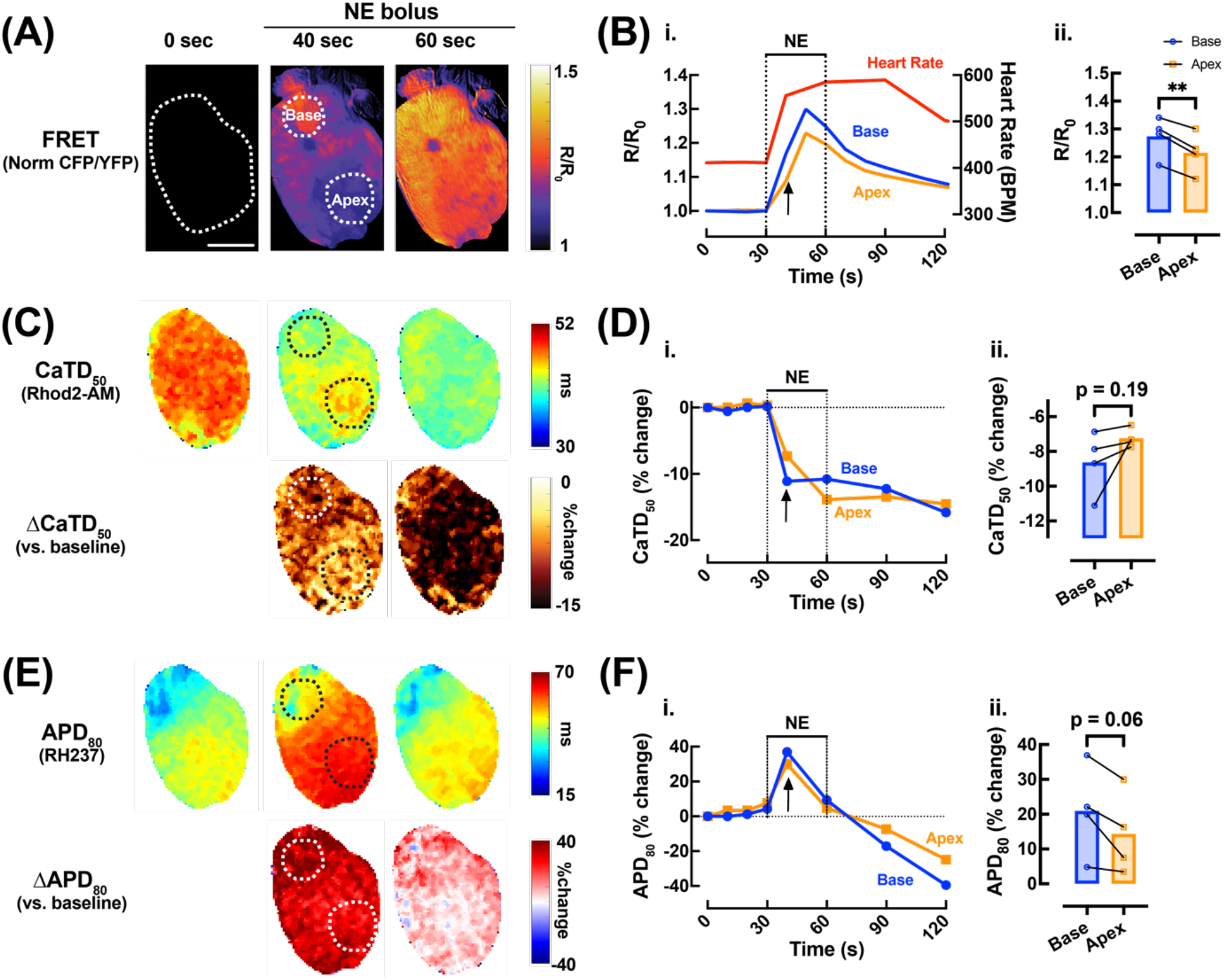
Effects β-adrenergic stimulation on cAMP responsiveness and Ca^2+^_i_ heterogeneity in the *CAMPER* mouse heart. **(A)** Representative ΔFRET ratio images showing the spatio-temporal kinetics of cAMP activity (scale bar = 3 mm) and **(B**_**i**._) example FRET ratio trace and **(B**_**ii**._) mean scatter plots at time = 40 sec (black arrow) in response to bolus of 0.5 μM NE from different regions of the heart (blue = right ventricular (RV) base; orange = apex). **(C)** Representative CaTD_50_ and **(E)** APD_80_ maps from the heart in **(A)** (top) with change in CaTD_50_ and APD_80_ vs. baseline (ΔCaTD_50_ and ΔAPD_80_) (bottom). **(D**_**i**._) Example CaTD_50_ and **(F**_**i**._) APD_80_ percentage change trace over time and **(D**_**ii**._ **and F**_**ii**._) mean scatter plots from RV base (blue) and apex (orange) following bolus of 0.5 μM NE. Paired t-test, **p < 0.01, N = 4 hearts).

### Real-time atrial cAMP and V_m_-Ca^2+^_i_ response following β-AR stimulation

To assess the capability of our optical imaging platform in visualizing pacemaker and atrial cAMP activity and V_m_-Ca^2+^_i_ responses, Langendorff-perfused *CAMPER* hearts were positioned in the posterior view and the ventricles were covered by a piece of flexible black plastic sheet to minimize ventricular light scattering to the atrial region (Fig. 5A). Representative V_m_ and Ca^2+^_i_ traces recorded from the atria before and after an acute bolus of NE (1.5 μM) are shown in Fig. 5B. Corresponding cAMP FRET and V_m_ activation maps recorded after an acute bolus of NE are shown in Fig. 5C and 5D. The earliest site of activation within the intercaval region, identified from the V_m_ activation map, was used to functionally identify the leading pacemaker (sino-atrial node, SAN) in Fig. 5D. This region was colocalized with the cAMP FRET ratio image (Fig. 5C) to determine the cAMP response within the pacemaking region. The cAMP signal from the SAN region along with heart rate is plotted over time in Fig. 5E.

**Figure 5.**
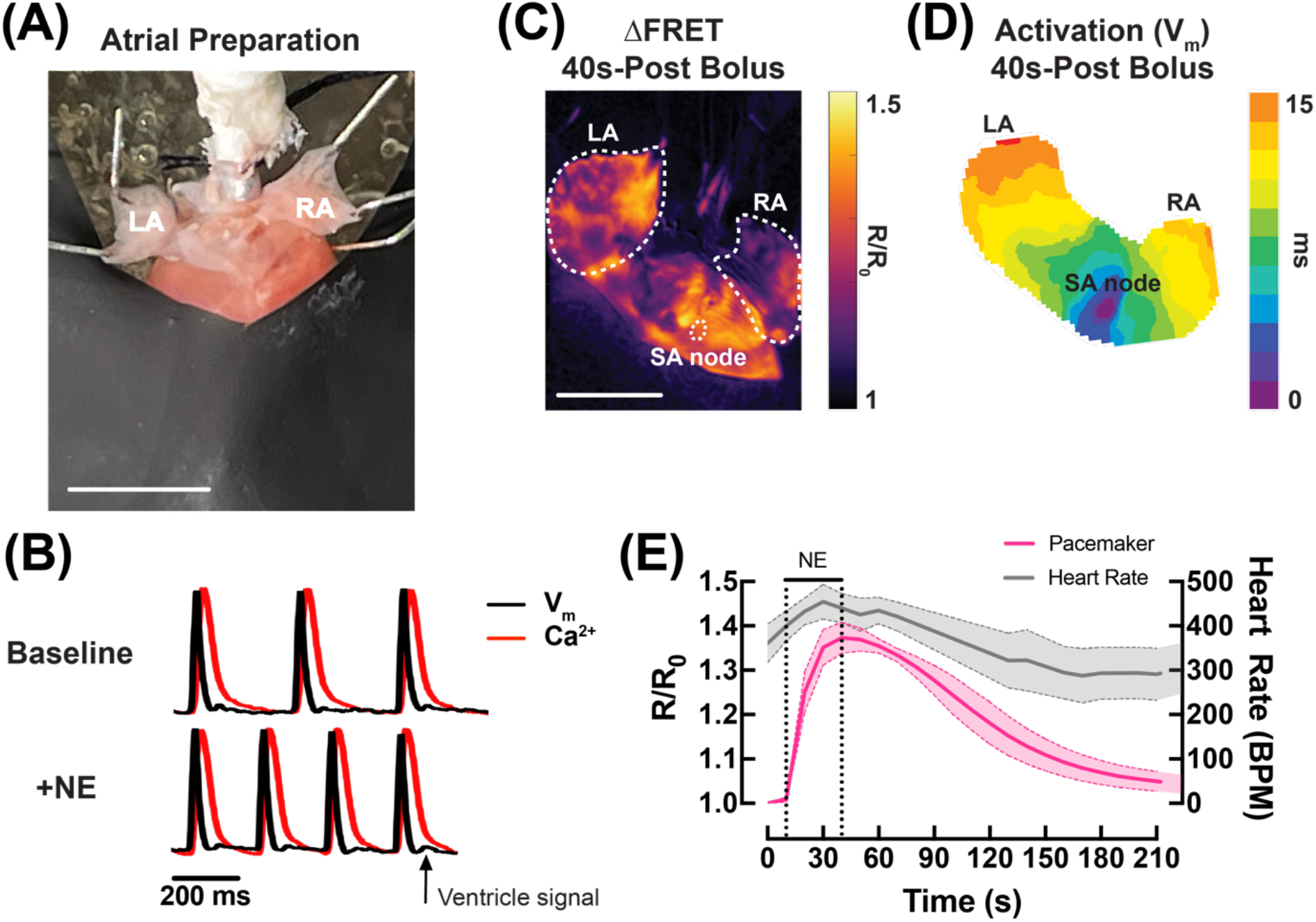
Atrial cAMP and V_m_-Ca^2+^_i_ response following β-adrenergic stimulation. **(A)** Example atrial preparation for optical mapping in a Langendorff-perfused *CAMPER* heart. Left atrium = LA, right atrium = RA. Scale bar = 3 mm. **(B)** Representative V_m_ and Ca^2+^ optical traces in the atria before and after a bolus of 1.5 μM NE. Black arrow indicates slight ventriclar signal bleed through due to light scattering. **(C)** Representative cAMP map and **(D)** V_m_ activation map in the atria after an acute bolus of 1.5 μM NE. Scale bar = 3 mm. **(E)** Averaged (mean = solid lines; SEM = shadow) temporal ΔFRET ratio traces in the sinoatrial node (SAN) with heart rate during bolus of 1.5 μM NE. N = 3 hearts.

### Whole-heart cAMP activity in response to sympathetic nerve stimulation

To assess whether the FRE-DOM system is capable of recording physiological levels of sympathetic activity (i.e., nerve-released NE), we compared whole-heart cAMP responses to exogenous NE versus neural stimulation. An acute bolus of NE (0.5 μM) was compared to chemically induced sympathetic nerve stimulation (SNS). An example spatio-temporal cAMP response was assessed from the left atrium (LA), RV base and apex of the heart after NE bolus. As can be observed in Fig. 6A, exogenous stimulation with NE produced relatively similar cAMP magnitude and kinetics in the RV base vs. apex, but lower in LA. Chemical SNS with a bolus of tyramine (20 μM, tyramine causes release of NE from nerve terminals) to the mouse heart was performed and the cAMP response is shown in Fig. 6B. Chemical SNS showed a higher cAMP response in the LA compared to the RV base and apex, suggesting differences in NE release between the atrial and ventricular regions.

**Figure 6.**
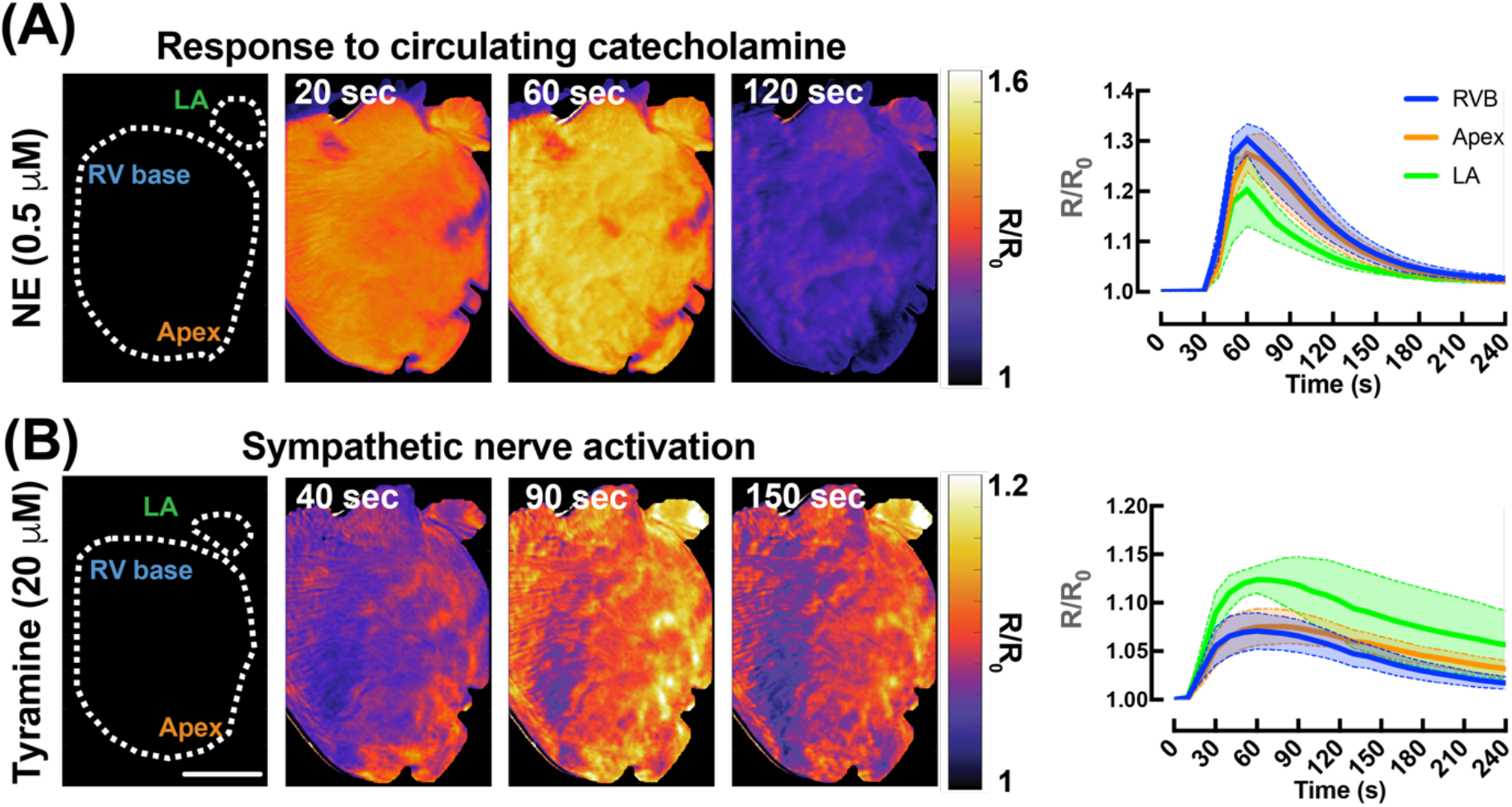
Effects of sympathetic activation on whole-heart cAMP activity. **(A)** Representative ΔFRET ratio images showing the spatio-temporal kinetics of cAMP activity (scale bar = 3 mm) in response to bolus of 0.5 μM NE or **(B)** 20 μM tyramine and averaged (mean = solid line; SEM = shadow) temporal FRET traces from different regions of the heart (green = LA; blue = RV base; orange = apex). N = 4 hearts.

## Discussion

In this study, we developed and validated an innovative multi-parametric imaging system to capture real-time, whole-heart interactions between adrenergic signaling pathways and the corresponding functional outcomes during sympathetic activation. To our knowledge, this is the first study to integrate simultaneous FRET imaging and dual optical mapping in an intact mouse heart. Key findings include robust cAMP activation across the both atrial and ventricular regions in response to β-AR stimulation, synchronized with changes in heart rate, action potential, and calcium transient responses. Additionally, our mapping system demonstrated flexibility for imaging different cardiac regions, responses to both exogenous and endogenous catecholamines, and integrating alternative calcium dyes to improve spectral separation and to support alternative fluorophore combinations. Building on these advancements, our approach offers the potential to expand multi-parametric imaging toward other nodes of the β-AR signaling pathway. This is particularly relevant given recent advances in genetically encoded fluorescent biosensors, such as GPCR activation-based biosensors, which are widely used in a variety of neuroscience applications and could be used in cardiac signaling with similar precision^30,31^. Such tools could significantly expand the scope of multi-parametric imaging, providing critical insights into the molecular mechanisms underlying cardiac dysfunction.

Our earlier research utilized a modified conventional cardiac optical mapping system to simultaneously measure cAMP signaling and V_m_ in the intact mouse heart. This study provided critical insights, revealing sex-dependent regional heterogeneity in cAMP breakdown and associated changes in APD and repolarization^19^. Such findings underscore the complexity of signaling pathways in cardiac function and their contribution to arrhythmogenic processes. While that study was the first to successfully image molecular signaling and electrophysiological responses simultaneously at the whole-heart level, it did not address calcium dynamics. In the present study, we found regional variation in cAMP activity (Fig. 4B_ii._), which corresponded to functional differences in both CaTD_50_ and APD_80_ between the RV base and apex. Although these electrophysiological differences were not statistically significant, we observed a trend towards greater shortening of CaTD_50_ and prolongation of APD_80_ in the RV base of the heart compared with the apex (Fig. 4D _ii._ and 4F _ii._). These results suggest elevated cAMP-PKA-mediated phosphorylation of phospholamban and L-type calcium channels (LTCCs) in the RV base, potentially enhancing sarcoplasmic reticulum (SR) calcium uptake and increase the inward L-type calcium current, respectively. As these results are from mixed-sex animals, more significant sex-dependent regional heterogeneity may emerge in future sex-separated analyses. cAMP and calcium mishandling are well-established contributors to arrhythmogenesis, particularly during pathological conditions such as myocardial infarction (MI) and heart failure. Elevated cAMP levels, while essential for regulating cardiac contractility and heart rate under physiological conditions, can lead to detrimental effects in disease states. These include cytosolic calcium overload, which promotes triggered activity through delayed afterdepolarizations, facilitates conditions for reentrant arrhythmias, and disrupts intercellular coupling. Collectively, these alterations contribute to the initiation and maintenance of arrhythmias^32^.

Cardiac optical mapping of intracellular calcium dynamics and V_m_ is standard for studying excitation-contraction coupling and cardiac rhythms. However, it provides limited insight into upstream molecular drivers. In recent years, targeted biosensors have been used to monitor compartmentalized cAMP signaling in isolated cardiomyocytes, revealing the impact of receptor-microdomain disruptions in disease states^33^. However, more recent studies have begun to assess cardiomyocyte signaling events within the intact heart. For example, Jungen *et al*. demonstrated how parasympathetic innervation regulates ventricular arrhythmia susceptibility via reductions in ventricular cAMP and shortening of refractory periods^34^. Similarly, metabolic indicators, such as NADH, further highlight the need for integrative approaches. Variability in NADH fluctuations under stress conditions underscores differences between to Langendorff-perfused hearts and their responses to hypoxia, ischemia, or increased workload^35^. These findings emphasize the utility of assessing signaling events in real-time in intact tissues and hearts.

Our group, along with others, are now pioneering efforts to combine traditional optical mapping with techniques that simultaneously reveal molecular signatures. This innovative approach bridges the gap between upstream molecular signaling and downstream electrophysiological outcomes, enabling a deeper understanding of the interplay between signaling, metabolism, excitation, and contraction in cardiac tissues. A notable advancement was recently reported by George *et al*., who used triple-parametric optical mapping to simultaneously measure Ca^2+^_i_, V_m_, and NADH, to investigate how metabolic state influences electrical and contractile behavior^36^. Their findings highlighted how changes in metabolic activity influence electrophysiological responses and calcium handling, shedding light on the mechanisms underlying arrhythmogenesis and myocardial dysfunction in these settings. Future research should continue building on these advancements, focusing on the integration of additional molecular markers, and applying these methods to disease models, to enhance the scope of cardiac optical mapping and its translational potential.

### Limitations

Mice expressing the cAMP biosensor *in vivo* were utilized in the study. However, mice have distinct differences in AP kinetics and ion channel expression compared to humans^37^, which may limit the translational relevance of the findings. Previous work from our group has characterized species-specific responses in AP kinetics to sympathetic activity, underscoring the need for caution when extrapolating mouse model results to human physiology^18^. The use of genetically encoded FRET biosensors, while powerful, also present potential limitations, including their influence on myocyte signaling networks. For example, cAMP buffering via binding to the Epac sensor may interfere with endogenous cAMP dynamics. Additionally, expression levels of the FRET biosensor may vary between individual hearts and affect quantitative comparisons. To minimize motion artifacts during optical imaging, hearts were perfused with blebbistatin, a myosin II inhibitor that suppresses contraction. However, blebbistatin may have secondary effects on electrical activity, metabolism, and coronary flow^27^. Finally, this study employed a novel multi-parametric imaging system based on a tandem-lens epifluorescence macroscope setup. While, this system offers substantial advantages in wide-field imaging, such as a 100-700 times brighter fluorescence image and improved light collection efficiency^21^, it lacks the spatial and depth resolution of traditional high-resolution microscopy systems. These constraints may limit the ability to capture fine structural details or subcellular signaling domains.

## Aknowledgement

We thank Brady Okura, Toshi Sakuraba, and Kenji Tsubokura from SciMedia, USA for technical support. We thank Gianna Domeny for maintaining, breeding and genotyping the cardiac-specific *CAMPER* mouse line.

## Funding

This work was supported by grants from the National Institutes of Health (NIH) R01 HL111600, HL170626 (C.M.R), NIH K99 HL171836 (J.L.C.), and NIH T32 GM144303 (L.R.M).

## Author Contributions

Conceptualization and supervision: C.M.R. Methodology: I.-J.E.L., J.L.C., C.M.R. Data generation: I.-J.E.L., J.L.C., L.R.M., L.W. Data analysis: I.-J.E.L., J.L.C., L.R.M. Writing of the manuscript: I.-J.E.L., C.M.R. Review & editing of the manuscript: I.-J. E. L., J.L.C., L.R.M., L.W., C.M.R.

## Competing Interests

Authors declare that they have no competing interests.

## Data and materials availability

All data, code, and materials are available from the authors upon request.

## Notes

### Competing Interest Statement

The authors have declared no competing interest.

## Reference

1. Calamera, G., Li, D., Ulsund, A.H., Kim, J.J., Neely, O.C., Moltzau, L.R., Bjørnerem, M., Paterson, D., Kim, C., Levy, F.O., and Andressen, K.W. (2019). FRET-based cyclic GMP biosensors measure low cGMP concentrations in cardiomyocytes and neurons. Communications Biology 2, 394. 10.1038/s42003-019-0641-x.

2. DiPilato, L.M., Cheng, X., and Zhang, J. (2004). Fluorescent indicators of cAMP and Epac activation reveal differential dynamics of cAMP signaling within discrete subcellular compartments. Proceedings of the National Academy of Sciences 101, 16513–16518. 10.1073/pnas.0405973101.

3. Surdo, N.C., Berrera, M., Koschinski, A., Brescia, M., Machado, M.R., Carr, C., Wright, P., Gorelik, J., Morotti, S., Grandi, E., et al. (2017). FRET biosensor uncovers cAMP nano-domains at β-adrenergic targets that dictate precise tuning of cardiac contractility. Nature Communications 8, 15031. 10.1038/ncomms15031.

4. Götz, K.R., and Nikolaev, V.O. (2013). Advances and Techniques to Measure cGMP in Intact Cardiomyocytes. In Guanylate Cyclase and Cyclic GMP: Methods and Protocols, T. Krieg, and R. Lukowski, eds. (Humana Press), pp. 121–129. 10.1007/978-1-62703-459-3_7.

5. Barbagallo, F., Xu, B., Reddy, G.R., West, T., Wang, Q., Fu, Q., Li, M., Shi, Q., Ginsburg, K.S., Ferrier, W., et al. (2016). Genetically Encoded Biosensors Reveal PKA Hyperphosphorylation on the Myofilaments in Rabbit Heart Failure. Circulation Research 119, 931–943. 10.1161/CIRCRESAHA.116.308964.

6. Liu, S., Zhang, J., and Xiang, Y.K. (2011). FRET-based direct detection of dynamic protein kinase A activity on the sarcoplasmic reticulum in cardiomyocytes. Biochemical and Biophysical Research Communications 404, 581–586. 10.1016/j.bbrc.2010.11.116.

7. Erickson, J.R., Patel, R., Ferguson, A., Bossuyt, J., and Bers, D.M. (2011). Fluorescence Resonance Energy Transfer–Based Sensor Camui Provides New Insight Into Mechanisms of Calcium/Calmodulin-Dependent Protein Kinase II Activation in Intact Cardiomyocytes. Circulation Research 109, 729–738. 10.1161/CIRCRESAHA.111.247148.

8. Stroik, D.R., Ceholski, D.K., Bidwell, P.A., Mleczko, J., Thanel, P.F., Kamdar, F., Autry, J.M., Cornea, R.L., and Thomas, D.D. (2020). Viral expression of a SERCA2a-activating PLB mutant improves calcium cycling and synchronicity in dilated cardiomyopathic hiPSC-CMs. Journal of Molecular and Cellular Cardiology 138, 59–65. 10.1016/j.yjmcc.2019.11.147.

9. Xie, Y., Sato, D., Garfinkel, A., Qu, Z., and Weiss, J.N. (2010). So Little Source, So Much Sink: Requirements for Afterdepolarizations to Propagate in Tissue. Biophysical Journal 99, 1408–1415. 10.1016/j.bpj.2010.06.042.

10. Bird, S.D., Doevendans, P.A., van Rooijen, M.A., Brutel de la Riviere, A., Hassink, R.J., Passier, R., and Mummery, C.L. (2003). The human adult cardiomyocyte phenotype. Cardiovascular Research 58, 423–434. 10.1016/s0008-6363(03)00253-0.

11. Mantravadi, R., Gabris, B., Liu, T., Choi, B.-R., de Groat, W.C., Ng, G.A., and Salama, G. (2007). Autonomic Nerve Stimulation Reverses Ventricular Repolarization Sequence in Rabbit Hearts. Circulation Research 100, e72–e80. 10.1161/01.RES.0000264101.06417.33.

12. Efimov, I.R., Nikolski, V.P., and Salama, G. (2004). Optical Imaging of the Heart. Circulation Research 95, 21–33. 10.1161/01.RES.0000130529.18016.35.

13. Avula, U.M.R., Abrams, J., Katchman, A., Zakharov, S., Mironov, S., Bayne, J., Roybal, D., Gorti, A., Yang, L., Iyer, V., et al. (2019). Heterogeneity of the action potential duration is required for sustained atrial fibrillation. JCI Insight 4. 10.1172/jci.insight.128765.

14. Wang, L., Myles, R.C., Lee, I.-J., Bers, D.M., and Ripplinger, C.M. (2021). Role of Reduced Sarco-Endoplasmic Reticulum Ca2+-ATPase Function on Sarcoplasmic Reticulum Ca2+ Alternans in the Intact Rabbit Heart. Frontiers in Physiology 12. 10.3389/fphys.2021.656516.

15. Rajendran, P.S., Challis, R.C., Fowlkes, C.C., Hanna, P., Tompkins, J.D., Jordan, M.C., Hiyari, S., Gabris-Weber, B.A., Greenbaum, A., Chan, K.Y., et al. (2019). Identification of peripheral neural circuits that regulate heart rate using optogenetic and viral vector strategies. Nature Communications 10, 1944. 10.1038/s41467-019-09770-1.

16. Zhu, C., Rajendran, P.S., Hanna, P., Efimov, I.R., Salama, G., Fowlkes, C.C., and Shivkumar, K. (2022). High-resolution structure-function mapping of intact hearts reveals altered sympathetic control of infarct border zones. JCI Insight 7. 10.1172/jci.insight.153913.

17. Muntean, B.S., Zucca, S., MacMullen, C.M., Dao, M.T., Johnston, C., Iwamoto, H., Blakely, R.D., Davis, R.L., and Martemyanov, K.A. (2018). Interrogating the Spatiotemporal Landscape of Neuromodulatory GPCR Signaling by Real-Time Imaging of cAMP in Intact Neurons and Circuits. Cell Reports 24, 1081–1084. 10.1016/j.celrep.2018.07.031.

18. Wang, L., Morotti, S., Tapa, S., Francis Stuart, S.D., Jiang, Y., Wang, Z., Myles, R.C., Brack, K.E., Ng, G.A., Bers, D.M., et al. (2019). Different paths, same destination: divergent action potential responses produce conserved cardiac fight-or-flight response in mouse and rabbit hearts. The Journal of Physiology 597, 3867–3883. 10.1113/JP278016.

19. Caldwell, J.L., Lee, I.-J., Ngo, L., Wang, L., Bahriz, S., Xu, B., Bers, D.M., Navedo, M.F., Bossuyt, J., Xiang, Y.K., and Ripplinger, C.M. (2023). Whole-heart multiparametric optical imaging reveals sex-dependent heterogeneity in cAMP signaling and repolarization kinetics. Science Advances 9, eadd5799. 10.1126/sciadv.add5799.

20. Muntean, B.S., Zucca, S., MacMullen, C.M., Dao, M.T., Johnston, C., Iwamoto, H., Blakely, R.D., Davis, R.L., and Martemyanov, K.A. (2018). Interrogating the Spatiotemporal Landscape of Neuromodulatory GPCR Signaling by Real-Time Imaging of cAMP in Intact Neurons and Circuits. Cell Reports 22, 255–268. 10.1016/j.celrep.2017.12.022.

21. Ratzlaff, E.H., and Grinvald, A. (1991). A tandem-lens epifluorescence macroscope: Hundred-fold brightness advantage for wide-field imaging. Journal of Neuroscience Methods 36, 127–137. 10.1016/0165-0270(91)90038-2.

22. Hallum, L.E., Chen, S.C., Cloherty, S.L., Morley, J.W., Suaning, G.J., and Lovell, N.H. (2006). Functional Optical Imaging of Intrinsic Signals in Cerebral Cortex. In Wiley Encyclopedia of Biomedical Engineering. 10.1002/9780471740360.ebs1588.

23. Couto, J., Musall, S., Sun, X.R., Khanal, A., Gluf, S., Saxena, S., Kinsella, I., Abe, T., Cunningham, J.P., Paninski, L., and Churchland, A.K. (2021). Chronic, cortex-wide imaging of specific cell populations during behavior. Nature Protocols 16, 3241–3263. 10.1038/s41596-021-00527-z.

24. Wang, Z., Wang, L., Tapa, S., Pinkerton, K.E., Chen, C.-Y., and Ripplinger, C.M. (2018). Exposure to Secondhand Smoke and Arrhythmogenic Cardiac Alternans in a Mouse Model. Environmental Health Perspectives 126, 127001. 10.1289/EHP3664.

25. Wang, Z., Tapa, S., Francis Stuart, S.D., Wang, L., Bossuyt, J., Delisle, B.P., and Ripplinger, C.M. (2020). Aging Disrupts Normal Time-of-Day Variation in Cardiac Electrophysiology. Circulation: Arrhythmia and Electrophysiology 13, e008093. 10.1161/CIRCEP.119.008093.

26. Tapa, S., Wang, L., Francis Stuart, S.D., Wang, Z., Jiang, Y., Habecker, B.A., and Ripplinger, C.M. (2020). Adrenergic supersensitivity and impaired neural control of cardiac electrophysiology following regional cardiac sympathetic nerve loss. Scientific Reports 10, 18801. 10.1038/s41598-020-75903-y.

27. Swift, L.M., Kay, M.W., Ripplinger, C.M., and Posnack, N.G. (2021). Stop the beat to see the rhythm: excitation-contraction uncoupling in cardiac research. American Journal of Physiology-Heart and Circulatory Physiology 321, H1005–H1013. 10.1152/ajpheart.00477.2021.

28. O’Shea, C., Holmes, A.P., Yu, T.Y., Winter, J., Wells, S.P., Correia, J., Boukens, B.J., De Groot, J.R., Chu, G.S., Li, X., et al. (2019). ElectroMap: High-throughput open-source software for analysis and mapping of cardiac electrophysiology. Scientific Reports 9, 1389. 10.1038/s41598-018-38263-2.

29. Ripplinger, C.M., Glukhov, A.V., Kay, M.W., Boukens, B.J., Chiamvimonvat, N., Delisle, B.P., Fabritz, L., Hund, T.J., Knollmann, B.C., Li, N., et al. (2022). Guidelines for assessment of cardiac electrophysiology and arrhythmias in small animals. American Journal of Physiology-Heart and Circulatory Physiology 323, H1137–H1166. 10.1152/ajpheart.00439.2022.

30. Feng, J., Zhang, C., Lischinsky, J.E., Jing, M., Zhou, J., Wang, H., Zhang, Y., Dong, A., Wu, Z., Wu, H., et al. (2019). A Genetically Encoded Fluorescent Sensor for Rapid and Specific In Vivo Detection of Norepinephrine. Neuron 102, 745-761.e748. 10.1016/j.neuron.2019.02.037.

31. Feng, J., Dong, H., Lischinsky, J.E., Zhou, J., Deng, F., Zhuang, C., Miao, X., Wang, H., Li, G., Cai, R., et al. (2024). Monitoring norepinephrine release in vivo using next-generation GRABNE sensors. Neuron 112, 1930-1942.e1936. 10.1016/j.neuron.2024.03.001.

32. Lubbe, W.F., Podzuweit, T., and Opie, L.H. (1992). Potential arrhythmogenic role of cyclic adenosine monophosphate (AMP) and cytosolic calcium overload: Implications for prophylactic effects of beta-blockers in myocardial infarction and proarrhythmic effects of phosphodiesterase inhibitors. Journal of the American College of Cardiology 19, 1622–1633. 10.1016/0735-1097(92)90629-2.

33. Sprenger, J.U., Perera, R.K., Steinbrecher, J.H., Lehnart, S.E., Maier, L.S., Hasenfuss, G., and Nikolaev, V.O. (2015). In vivo model with targeted cAMP biosensor reveals changes in receptor–microdomain communication in cardiac disease. Nature Communications 6, 6965. 10.1038/ncomms7965.

34. Jungen, C., Scherschel, K., Eickholt, C., Kuklik, P., Klatt, N., Bork, N., Salzbrunn, T., Alken, F., Angendohr, S., Klene, C., et al. (2017). Disruption of cardiac cholinergic neurons enhances susceptibility to ventricular arrhythmias. Nature Communications 8, 14155. 10.1038/ncomms14155.

35. Wengrowski, A.M., Kuzmiak-Glancy, S., Jaimes, R., and Kay, M.W. (2014). NADH changes during hypoxia, ischemia, and increased work differ between isolated heart preparations. American Journal of Physiology-Heart and Circulatory Physiology 306, H529–H537. 10.1152/ajpheart.00696.2013.

36. George, S.A., Lin, Z., and Efimov, I.R. (2022). Simultaneous triple-parametric optical mapping of transmembrane potential, intracellular calcium and NADH for cardiac physiology assessment. Commun Biol 5, 319. 10.1038/s42003-022-03279-y.

37. Bartos, D.C., Grandi, E., and Ripplinger, C.M. (2015). Ion Channels in the Heart. Compr Physiol 5, 1423–1464. 10.1002/cphy.c140069.

